# Aliquoting of isobaric labeling reagents for low concentration and single cell proteomics samples

**DOI:** 10.1101/2021.06.23.449560

**Authors:** Yuting Yuan, Benjamin C. Orsburn

## Abstract

The introduction of isobaric tagging reagents enabled more accurate, high-throughput quantitative proteomics by enabling multiple samples to be multiplexed. One drawback of these workflows is the relative expense of the proprietary isobaric reagents, which is often only second to the expense of the instruments themselves. These highly reactive chemical tags are only commercially available in relatively large aliquots compared to the typical amounts of peptides analyzed in proteomic workflows today. Excess reagents are typically disposed of following a single labeling experiment or those performed within a few days of opening a new kit. We present a simple procedure to aliquot commercial isobaric tagging reagents and demonstrate the successful and high efficiency labeling of multiple samples over a period of six months. The samples presented herein were selected as the most diverse ones labeled by prepared aliquots from a single labeling reagent kit over this period. We observe comparable labeling efficiency from 100 microgram to 100 picograms of peptide when labeling samples from both human digest standards, cancer cell lines prepared in-house and from cells directly obtained from human organ donors, despite differences in cell type, lysis, and digestion procedures. No labeling experiment of whole human proteomics samples achieved less than 92% labeling efficiency over this period. When preparing phosphoproteomic samples 6 months after the date of the aliquoting procedure, we observed a decrease in labeling efficiency to approximately 86%, indicating the end of the useful lifetime of aliquots prepared in this manner. Over this period, we have effectively reduced the reagent costs of each experiment to less than 10% of the predicted costs when following the manufacturer instructions for use and disposal. While aliquoting of reagents can be performed by hand, we provide a complete template for automatic aliquoting using an affordable liquid handling robot, including plans for 3D printing of two parts we have found useful for streamlining this procedure.

**Abstract Graphic:** 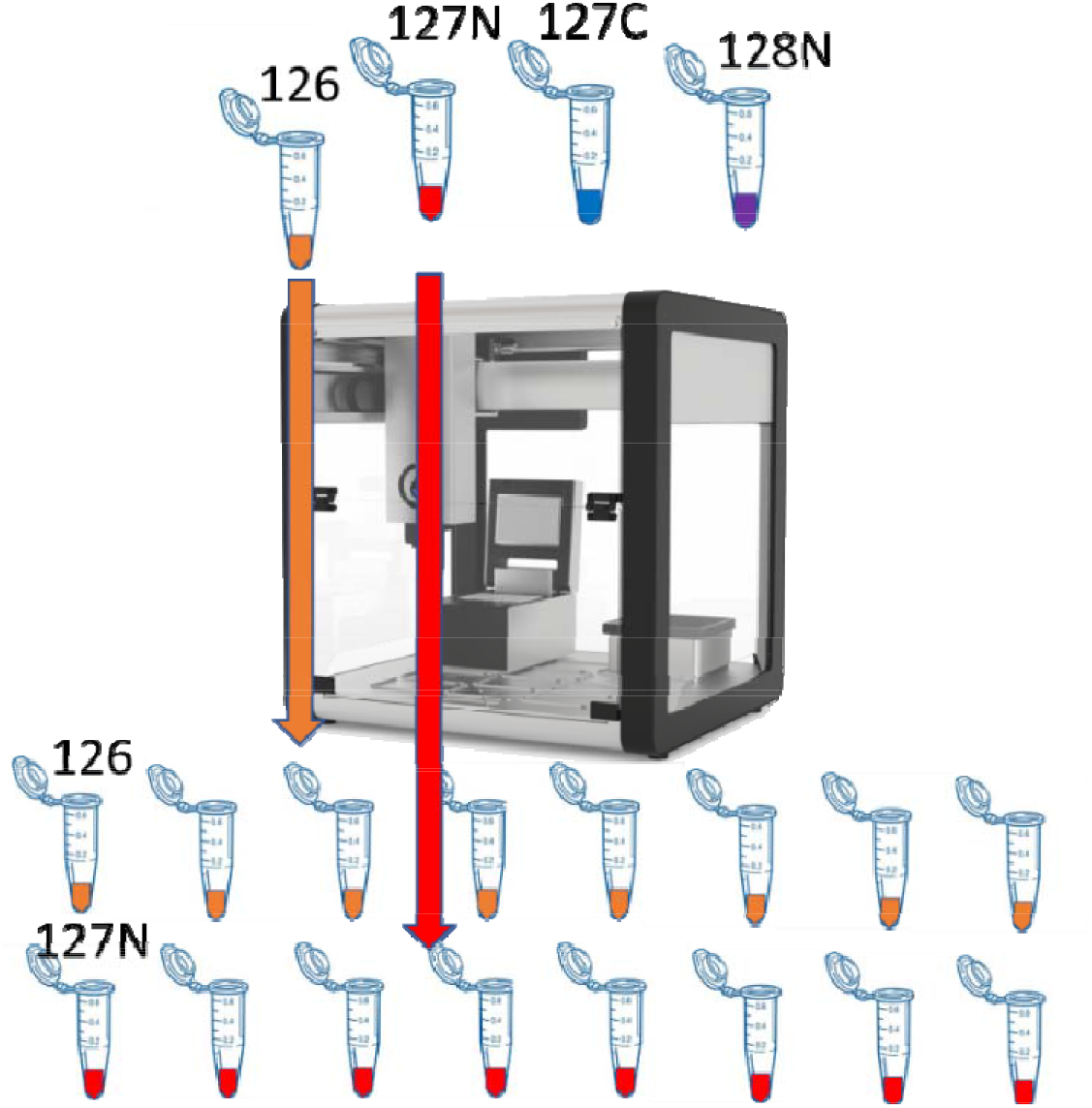

## Introduction

The use of high performance liquid chromatography coupled to tandem mass spectrometry (LCMS) for the identification and quantification of enzymatically digested proteins in biological samples is rapidly becoming a routine and complementary method to genomic analyses.^1^ Although different at the basic technological level, both genomic and proteomics technologies benefit from the use of multiplexing as a primary cost and time reduction mechanism.^2,3^ In addition, recent methods using tagged peptides in specifically altered proportions are used to BOOST some peptide signals, further increasing their usefulness in the lab.^4^ In proteomics, enzymatically digested peptides are labeled with isobaric chemical tags (IBT). Following tagging, the labeled peptide mixtures are combined. The resulting mixture of labeled peptides are indistinguishable at nearly every level due to the swapping of heavy isotopic labels between a reporter and balance region on the tags. Upon MS/MS fragmentation, the reporter and balance regions of the tag separate, producing a low mass fragment ion that can be used for relative protein quantifications of the individual samples. Technologies such as the Tandem mass tag (TMT*) and isobaric tag for relative protein quantification (iTRAQ*) are used in nearly every proteomics lab today due to the ability to obtain quantitative data of up to 16 proteomics samples at nearly the same level of protein coverage as a single samples using label free relative quantification.^5,6^ IBT reagents are the exclusive trademarked properties of a single manufacturer and, while these reagents are distributed by multiple vendors in different formats, the costs per kit are often the largest daily expenditure in proteomics labs. Recent solutions have been described to help lower these costs, including reducing the relative concentration of labeling reagents to peptide samples compared to manufacturer recommendations.^7,8^

Today’s LCMS systems are largely faster and more sensitive than the instruments of the past. As such, today’s experiments routinely assess samples that were previously unthinkable. The most extreme example is likely the recent description of Single Cell Proteomics by Mass Spectrometry (SCoPE-MS) in which individual cells are lysed, digested and labeled.^9–11^ Other examples include the near routine descriptions of temporal proteomics,^12^ and the analysis of increasingly smaller samples removed from tissues by procedures such as microcapture laser dissection.^13^ The typical mammalian cell contains between 50 and 650 picograms of protein.^14^ When employing the manufacturer’s recommended labeling procedure of 10:1 reagent to even large cell such as a human hepatocyte, one aliquot of reagent can be used to label over 10,000 individual cells. When multiplexing is considered, this label efficiency may represent as many as 140,000 single cells. Using the standard two hour LCMS experiment time described in the SCOPE2 protocol, over 2 years of instrument time would be needed to fully utilize this single reagent kit (data not shown).

Core labs or certified research organizations (CROs) that predominantly focus on labeled proteomics experiments may be able to effectively utilize large reagent kits. For research labs like ours which use labeled proteomics as only one of many workflows, rapidly expiring kits (and waste many spare reagents/original reagents bought from vendors are not fully utilized and it is soon expired if not stored properly) may be a major expenditure, leading to the adoption of alternative protocols.^8^ Using kits that were opened for single experiments, we developed a simple and rapid aliquoting procedure could vastly expand the life and usefulness of individual reagent kits. Additional development of this procedure with inexpensive automation has allowed us to routinely prepare reagents in concentrations appropriate for single cell proteomics applications.

## Methods

### Development of liquid handling protocol

The OT-2 liquid handler robot was fitted with a P20 G2 20 μL digital pipettor (OpenTrons, Brooklyn, NY). Methods were developed through the OT2 graphical user interface (GUI) based off the “Simple Aliquot Protocol”. Adjustments were made to the method by visually monitoring the transfer of anhydrous acetonitrile. To minimize the exposure of the reagents to light, the main deck lights on the liquid handler were disabled through the GUI when possible and covered with lab labeling tape when these could not be disabled. The liquid handler deck was equipped with two “4-in-one” tube racks (OpenTrons). Due to the inability to use differential heights for sample pickup and distribution through the GUI, an additional tube rack was fabricated using 3D plans obtained from the manufacturer library. The height of the 1.5 mL receiving rack plate was reduced in width to 75% of its original dimension using the open source software Cura 17.0.2. (Ultimaker) The plans for the initial rack and the adjusted product are provided in the supplemental material. The rack was fabricated with 1.75 mm polylactic acid filament (HatchBox) using a modular table 3D printer (MOD-T).

### Reagent suspension and aliquoting

The TMTPro 16-plex IBT reagent kit in 0.5 μg aliquots was purchased from Thermo Fisher Scientific. Each aliquot was resuspended in 120 μL of anhydrous acetonitrile removed from a self-sealing container by gas tight 200 μL syringe. All reagents were pipetted into 1.5 mL Eppendorf tubes on dry ice. Aliquots of reagent were used for sample labeling of a serial dilution of K562 peptide digest standard (Promega) using all heavy N labeled channels that day following manufacturer instructions. Ten microliter aliquots were rapidly transferred from the OpenTrons OT-2 rack using the OpenTrons GUI and the TMT_aliquotting.json file provided in the supplemental material. Following the automated aliquoting procedure, tubes were transferred to an air and light tight container of dry ice, prior to vacuum centrifugation (SpeedVac, Fisher) to dryness. To minimize light exposure all aliquoting by the OT-2 system was performed in the dark. All lights on the system were disabled or covered with labeling tape. Samples were removed from the liquid handler and placed into dry ice using a red led light source. The semi-transparent lid of the SpeedVac was covered in a sheet of aluminum foil to reduce light exposure. The 10 μL aliquots were fully dried in approximately 10 minutes at room temperature, sealed under red light and transferred to −80◻C for long term storage.

### Preparation and labeling of human cancer digests

Standard cancer digest K562 and HeLa were obtained from Promega and Pierce, respectively. NCI-H-358 cells were obtained from ATCC and grown according to ATCC protocols. Cells were lysed in S-Trap (ProtiFi) buffer and processed to peptides on S-Trap mini columns following manufacturer’s instructions. Proteins were reduced and alkylated using dithiothreitol and iodoacetamide when appropriate for the experiment. For single cell experiments and for samples to be analyzed for the global analysis of post-translational modifications these steps were not employed. Cancer cell line peptides were labeled with TMT reagents either freshly prepared following the instructions from Pierce for the kits, or dried aliquots were resuspended in anhydrous acetonitrile prior to labeling and manufacturer protocols were likewise followed. Labeling reactions were quenched in freshly prepared hydroxylamine. Library peptides were dried and subjected to offline fractionation using high pH reversed phase spin columns (Pierce) or desalted with C-18 spin columns (Pierce) according to manufacturer instructions.

### Preparation and labeling of human hepatocytes

Based on the former lab work (1), cryoplateable LIVERPOOL primary human (male and female) hepatocytes were purchased from BioIVT (Baltimore, MD). Cells were thawed and plated at a density of 700,000 viable cells/well on a 6-well collagen-coated clear flat-bottom plates (Corning) with *InVitroGro* CP medium (BioIVT) according to the manufacturer’s protocol with a viability of ≥ 90%. Following adherence 2-4 hours, we removed the culture medium and fresh treatment-containing *InVitroGro* CP medium supplemented with 50 μM acetaminophen was added into the hepatocytes culture (the concentration could induce substantial hepatotoxicity according to the previous lab study), and with vehicle (0.1% DMSO) was added into the hepatocytes as negative controls. Each acetaminophen and DMSO incubation wells had biological triplicates. After 24 hours incubation, the treated hepatocytes were harvested through cell scraping and lysed by using the 1x cell lysis buffer (Cell Signaling Technologies) supplemented with 1x Halt Protease and Phosphatase inhibitor Cocktail (Thermo Fisher Scientific), 0.5 mM phenylmethylsulfony fluoride (Millipore Sigma) and 0.5% sodium dodecyl sulfate (SDS), followed by centrifugation (3,000g, 5min, 4◻). Protein concentration was quantified using Qubit Protein Assay Kit (Thermo Fisher Scientific).

Cell lysate was diluted to 1mg/mL in 1x cell lysis buffer (Cell Signaling Technologies), supplemented with 0.5% SDS. Dithiothreitol (DTT) was added to a final concentration of 10 mM, and the sample was incubated for 45 min at 50 ◻. Iodoacetamide was added to a final concentration of 55mM, and the samples were incubated for 20 min and protected from light. Protein samples were loaded onto the suspension-trapping filter-based approach (S-Trap) mini columns (ProtiFi) according to the manufacturer’s instructions. In brief, samples were washed with 90:10% methanol/50 mM ammonium bicarbonate. Samples were then digested with trypsin (1:10 trypsin/protein) for 1 hour at 47 ◻. The digested peptides were eluted using 50 mM ammonium bicarbonate. Two more elutions were made using 0.2% formic acid and 0.2 % formic acid in 50% acetonitrile. The three elutions were pooled together and vacuum-centrifuged to dry. The digested elutions samples were reconstituted with 0.1% trifluoroacetic acid (TFA) in LC-MS grade water and loaded onto the Pierce peptide desalting spin columns (Thermo Fisher Scientific) according to the manufacturer’s protocols. The desalted peptides samples were eluted with 50% acetonitrile and 0.1% TFA water solution and vacuum-centrifuged to dry. Samples were kept at −80◻ until labeling.

### Single cell digest optimization and labeling

NCI-H-358 cells were obtained from ATCC and grown according to ATCC recommendations. Following 4 passages, cells were released from the plating medium with 0.5% trypsin solution at 37◻C for 15 minutes. The rounded cells were transferred to 15 mL conical tubes and pelleted at 500 x *g* for 5 minutes. The trypsin was poured off and the cells were rinsed once in 0.1% BSA with 1% trypsin inhibitor in HBS buffer (Fisher) and resuspended in 2 mL of the same. The cells were sorted into 96 well low protein bind plates (Dionex) at the Johns Hopkins University single cell sorting core. The individual cells were lysed and digested following the mPOP procedure.^15^ Based on data from the FACS sorting, cells were estimated to contain 200 picograms of total protein. Labeling with TMT 133C was performed at an approximate 10:1 ratio based on this estimate.

### Phosphopeptide enrichment

A twenty microgram digest of NCI H-358 peptide digest was labeled as described and enriched for phosphopeptides using the Fe-NTA spin column kits (Pierce) according to the manufacturer protocol. The enriched samples were directly transferred to C-18 spin columns (Pierce) for desalting prior to LCMS analysis.

### LC-MS Analysis

Samples were injected on an EasyNLC 1200 (Proxeon) with 0.1% formic acid in water as buffer A and 80% acetonitrile 0.1% formic acid coupled to a TIMSTOF Flex (Bruker) using experiment specific gradients and chromatography conditions. For bulk proteomics, a PepMap C-18 3 cm 2 μm trap column was used for sample pre concentration and desalting, for lower injection samples, the trap was not employed to minimize sample loss. The analytical column used for these experiments was a C-18 1.7 μm particle size, 75 μm x 15 cm PepSep column coupled by “zero dead volume” union (Bruker) to a 20 μm CaptiveSpray emitter. For experiments performed on the TIMSTOF “True Single Cell” prototype system, a Bruker NanoElute system was used with an IonOpticks Aurora 25cm column with integrated emitter. All LC and MS settings are included in each respective file included in each individual file in the data upload.

### Evaluation of labeling efficiency

Proteomic samples prepared from freshly aliquoted IBT reagents, as well as proteomic samples labeled over a subsequent period of 6 months following aliquoting and drying were converted to MGF files using ProteoWizard MSConvert Version: 3.0.20310-0d96039e2 (developer build) using the default settings for instrument conversion settings. The resulting MGF files were processed in IMP-Proteome Discoverer 2.4^16^ using the MSAmanda 2.0 node ^17^ with a 30 ppm MS1 and 0.05 Da MS/MS tolerance. Static carbamidomethylation was used for samples which were reduced and alkylated and not utilized when not appropriate. Oxidation of methionine was used as a dynamic option in all analyses. The IBT reagents were set as dynamic modifications on lysine and all peptide N-termini. For the phosphopeptide enriched samples, dynamic modifications were included on serine and threonine. After subtracting for common laboratory contaminants, the labeling efficiency is calculated as:

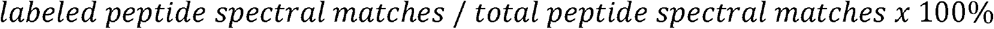

### Data Availability

All raw instrument files, MGF files used for analysis and Proteome Discoverer results have been uploaded to the FigShare public repository by project <u>11604</u>.

## Results and Discussion

### Labeling efficiency

Samples have been labeled with aliquoted TMT reagents for a series of different experiments, both for optimization of IBT experimental conditions and for biological samples. We have labeled cells from cancer cell lines HeLa, K562, and NCI-H-358 as well as from human donor hepatocytes. In all global experiments performed for the first 6 months, the PSM labeling efficiency exceeded 92% a number comparable to those described in other studies where reduction in TMT label usage has been described.^8,18^

**Figure 1.**
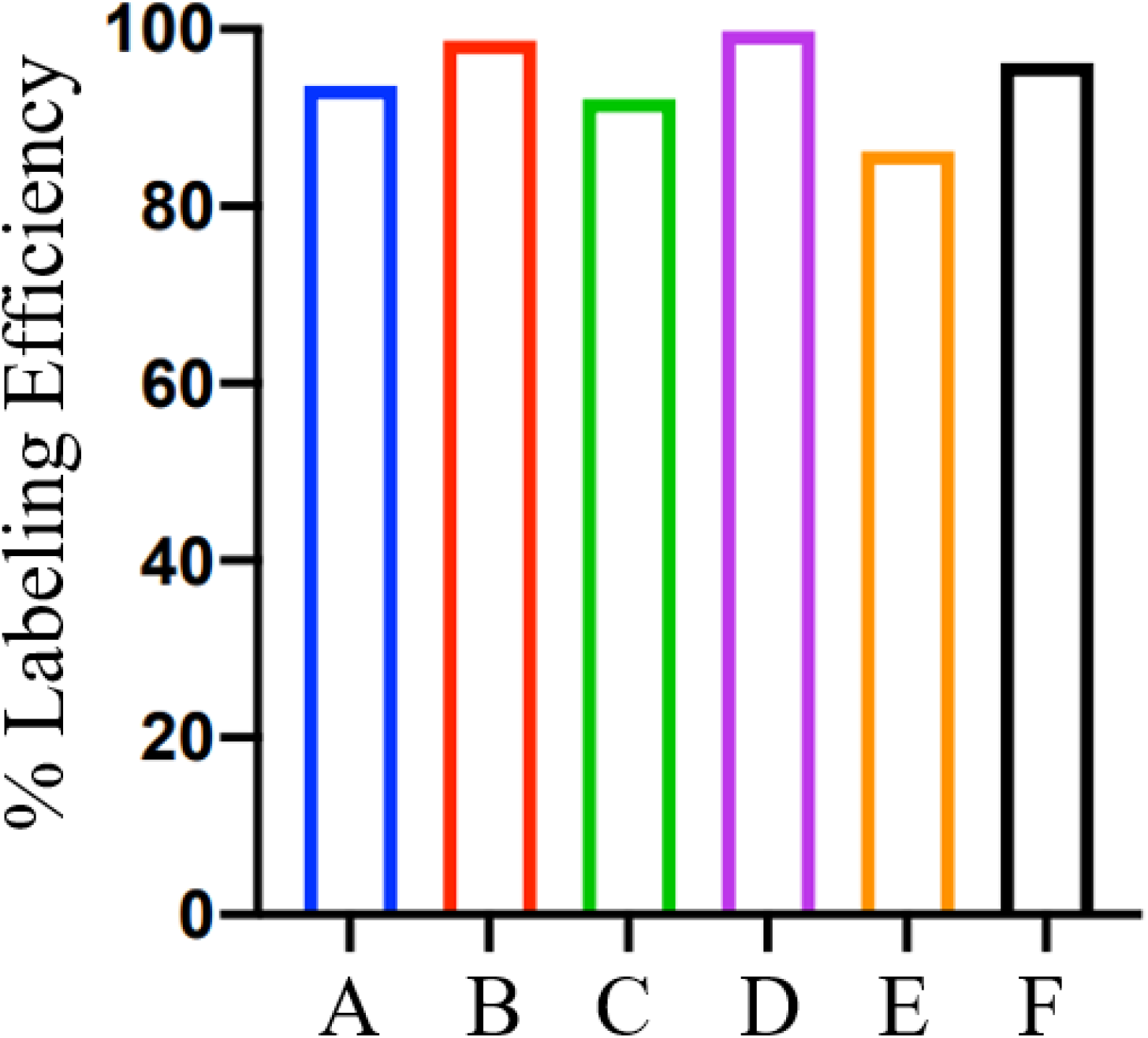
Calculated labeling efficiency of samples prepared over a 6 month period. (A) Labeled K562 digest, November 2020. (B) Liver hepatocytes from organ donor, December 2020. (C) K562 digest prepared February 2021. (D) H-358 cell lysate prepared March 2021. (E) H-358 lysate phosphopeptides prepared June 2021. (F) H-358 single cell preparation, May 2021.

### Relative cost savings

One TMTPro reagent kit was purchased in November of 2020 and opened on 11/17/2021. Following the manufacturer’s protocols allowing two weeks of storage of reagents in anhydrous acetonitrile, we would have performed 3 total labeling experiments with no alterations to our planned schedules. To date, we have performed a total of 11 separate labeling experiments utilizing aliquots from this initial kit, six of which have been described in this text, with the most recent use of a labeling reagent occurring on 5/15/2021. We often delay experiments to perform multiple labeling experiments when kits first arrive. Due to the spacing of experiments over the last 6 months, it is possible that we would have used a minimum of 3 reagent kits, and a maximum of 5. At an approximate $1,700 per kit, we have, to date, saved between $3,400 and $10,200 in reagent costs. At the date of this writing, reagent aliquots are still in storage from this kit for potential use.

## Conclusions

We have described a simple protocol that extends the practical usefulness of IBT reagent kits. The data presented herein demonstrates that a single kit may be aliquoted and stored with no obvious loss in sample labeling efficiency for at least 6 months from the date at which the kit is opened and aliquoted. It is worth noting that we have only performed this analysis using the recently described TMTPro 16-plex IBT reagent kit (Thermo Fisher Scientific). While we do not believe that this method should be exclusive to this IBT reagent kit, other reagents may require testing to determine whether these observations extend to other chemistries.

## References

(1) Rauniyar, N.; Yates 3rd, J. R. Isobaric Labeling-Based Relative Quantification in Shotgun Proteomics. J Proteome Res 2014, 13 (12), 5293–5309. https://doi.org/10.1021/pr500880b.

(2) Kircher, M.; Sawyer, S.; Meyer, M. Double Indexing Overcomes Inaccuracies in Multiplex Sequencing on the Illumina Platform. Nucleic Acids Res. 2012. https://doi.org/10.1093/nar/gkr771.

(3) Abbatiello, S. E.; Schilling, B.; Mani, D. R.; Zimmerman, L. J.; Hall, S. C.; MacLean, B.; Albertolle, M.; Allen, S.; Burgess, M.; Cusack, M. P.; Gosh, M.; Hedrick, V.; Held, J. M.; Inerowicz, H. D.; Jackson, A.; Keshishian, H.; Kinsinger, C. R.; Lyssand, J.; Makowski, L.; Mesri, M.; Rodriguez, H.; Rudnick, P.; Sadowski, P.; Sedransk, N.; Shaddox, K.; Skates, S. J.; Kuhn, E.; Smith, D.; Whiteaker, J. R.; Whitwell, C.; Zhang, S.; Borchers, C. H.; Fisher, S. J.; Gibson, B. W.; Liebler, D. C.; MacCoss, M. J.; Neubert, T. A.; Paulovich, A. G.; Regnier, F. E.; Tempst, P.; Carr, S. A. Large-Scale Interlaboratory Study to Develop, Analytically Validate and Apply Highly Multiplexed, Quantitative Peptide Assays to Measure Cancer-Relevant Proteins in Plasma. Mol. Cell. Proteomics 2015. https://doi.org/10.1074/mcp.M114.047050.

(4) Chua, X. Y.; Mensah, T.; Aballo, T.; Mackintosh, S. G.; Edmondson, R. D.; Salomon, A. R. Tandem Mass Tag Approach Utilizing Pervanadate BOOST Channels Delivers Deeper Quantitative Characterization of the Tyrosine Phosphoproteome. Mol. Cell. Proteomics 2020. https://doi.org/10.1074/mcp.TIR119.001865.

(5) Thompson, A.; Wölmer, N.; Koncarevic, S.; Selzer, S.; Böhm, G.; Legner, H.; Schmid, P.; Kienle, S.; Penning, P.; Höhle, C.; Berfelde, A.; Martinez-Pinna, R.; Farztdinov, V.; Jung, S.; Kuhn, K.; Pike, I. TMTpro: Design, Synthesis, and Initial Evaluation of a Proline-Based Isobaric 16-Plex Tandem Mass Tag Reagent Set. Anal. Chem. 2019. https://doi.org/10.1021/acs.analchem.9b04474.

(6) Wiese, S.; Reidegeld, K. A.; Meyer, H. E.; Warscheid, B. Protein Labeling by ITRAQ: A New Tool for Quantitative Mass Spectrometry in Proteome Research. Proteomics 2007. https://doi.org/10.1002/pmic.200600422.

(7) Navarrete-Perea, J.; Yu, Q.; Gygi, S. P.; Paulo, J. A. SL-TMT: A Streamlined Protocol for Quantitative (Phospho)Proteome Profiling Using TMT-SPS-MS3. J. Proteome Res. 2018. https://doi.org/10.1021/acs.jproteome.8b00217.

(8) Zecha, J.; Satpathy, S.; Kanashova, T.; Avanessian, S. C.; Kane, M. H.; Clauser, K. R.; Mertins, P.; Carr, S. A.; Kuster, B. TMT Labeling for the Masses: A Robust and Cost-Efficient, in-Solution Labeling Approach. Mol. Cell. Proteomics 2019. https://doi.org/10.1074/mcp.TIR119.001385.

(9) Specht, H.; Emmott, E.; Petelski, A. A.; Gray Huffman, R.; Perlman, D. H.; Serra, M.; Kharchenko, P.; Koller, A.; Slavov, N. Single-Cell Mass-Spectrometry Quantifies the Emergence of Macrophage Heterogeneity. bioRxiv. 2019. https://doi.org/10.1101/665307.

(10) Budnik, B.; Levy, E.; Harmange, G.; Slavov, N. SCoPE-MS: Mass Spectrometry of Single Mammalian Cells Quantifies Proteome Heterogeneity during Cell Differentiation. Genome Biol. 2018. https://doi.org/10.1186/s13059-018-1547-5.

(11) Slavov, N. Unpicking the Proteome in Single Cells. Science. 2020. https://doi.org/10.1126/science.aaz6695.

(12) Santana-Codina, N.; Chandhoke, A. S.; Yu, Q.; Małachowska, B.; Kuljanin, M.; Gikandi, A.; Stańczak, M.; Gableske, S.; Jedrychowski, M. P.; Scott, D. A.; Aguirre, A. J.; Fendler, W.; Gray, N. S.; Mancias, J. D. Defining and Targeting Adaptations to Oncogenic KRASG12C Inhibition Using Quantitative Temporal Proteomics. Cell Rep. 2020. https://doi.org/10.1016/j.celrep.2020.03.021.

(13) Kelly, R.; Zhu, Y.; Liang, Y.; Cong, Y.; Piehowski, P.; Dou, M.; Zhao, R.; Qian, W.-J.; Burnum-Johnson, K.; Ansong, C. Single Cell Proteome Mapping of Tissue Heterogeneity Using Microfluidic Nanodroplet Sample Processing and Ultrasensitive LC-MS. J. Biomol. Tech. 2019, 30 (Suppl), S61–S61.

(14) Wiśniewski, J. R.; Hein, M. Y.; Cox, J.; Mann, M. A “Proteomic Ruler” for Protein Copy Number and Concentration Estimation without Spike-in Standards. Mol. Cell. Proteomics 2014. https://doi.org/10.1074/mcp.M113.037309.

(15) Specht, H.; Harmange, G.; Perlman, D. H.; Emmott, E.; Niziolek, Z.; Budnik, B.; Slavov, N. Automated Sample Preparation for High-Throughput Single-Cell Proteomics. bioRxiv 2018. https://doi.org/10.1101/399774.

(16) Orsburn, B. C. Proteome Discoverer—A Community Enhanced Data Processing Suite for Protein Informatics. Proteomes 2021, 9 (1). https://doi.org/10.3390/proteomes9010015.

(17) Dorfer, V.; Pichler, P.; Stranzl, T.; Stadlmann, J.; Taus, T.; Winkler, S.; Mechtler, K. MS Amanda, a Universal Identification Algorithm Optimized for High Accuracy Tandem Mass Spectra. J. Proteome Res. 2014. https://doi.org/10.1021/pr500202e.

(18) Navarrete-Perea, J.; Yu, Q.; Gygi, S. P.; Paulo, J. A. Streamlined Tandem Mass Tag (SL-TMT) Protocol: An Efficient Strategy for Quantitative (Phospho)Proteome Profiling Using Tandem Mass Tag-Synchronous Precursor Selection-MS3. J. Proteome Res. 2018. https://doi.org/10.1021/acs.jproteome.8b00217.

